# Sensitive detection of protein ubiquitylation using a protein-fragment complementation assay

**DOI:** 10.1101/791897

**Authors:** Marie Le Boulch, Audrey Brossard, Gaëlle Le Dez, Gwenaël Rabut

## Abstract

Ubiquitylation is a reversible post-translational protein modification that regulates a multitude of cellular processes. Detection of ubiquitylated proteins is often challenging, because of their low abundance. Here, we present NUbiCA, a sensitive protein-fragment complementation assay to facilitate the monitoring of ubiquitylation events in cultured cells and model organisms. Using yeast as a model system, we demonstrate that NUbiCA enables to accurately monitor mono- and poly-ubiquitylation of proteins expressed at endogenous levels. We also show that it can be applied to decipher ubiquitin chain linkages. We used NUbiCA to investigate the ubiquitylation of the low abundance centromeric histone Cse4, and found that it is ubiquitylated during S-phase. Finally, we assembled a genome wide collection of yeast strains ready to investigate the ubiquitylation of proteins with this new assay. This resource will facilitate the analysis of local or transient ubiquitylation events that are difficult to detect with current methods.

**Summary statement:** We describe a sensitive protein-fragment complementation assay to facilitate the monitoring of ubiquitylation events that take place in cultured cells or model organisms.

## INTRODUCTION

Ubiquitylation is a prevalent posttranslational protein modification that plays a central role in the cell. It controls the homeostasis, turnover and activity of myriads of proteins. Defects in ubiquitylation are implicated in the etiology of numerous human diseases, including infection, neurodegenerative disorders, and cancers (Popovic et al., 2014). Ubiquitylation is catalyzed by ubiquitin conjugating enzymes (E2s) and ubiquitin ligases (E3s) that act in concert to covalently attach one or multiple ubiquitin moieties onto their substrate proteins, generally on lysine residues. E2s and E3s can also target the 7 lysine residues (K6, K11, K27, K29, K33, K48 and K63) and the amino terminus of ubiquitin to assemble various types of poly-ubiquitin chains. Depending on their topology, these ubiquitin chains act as distinct molecular signals that can have different consequences for the ubiquitylated proteins (Komander and Rape, 2012; Yau and Rape, 2016). For instance, K48- and K11-linked ubiquitin chains typically target the modified protein for proteasomal degradation, while K63-linked chains are associated with lysosomal degradation or non proteolytic regulatory mechanisms.

The elucidation of the functions and mechanisms of ubiquitylation demands sensitive tools to identify ubiquitylated proteins, monitor their modification and decipher ubiquitin linkages. Advances in mass spectrometry techniques and enrichment strategies now enable the identification of thousands of ubiquitylated proteins in cell extracts (Bennett et al., 2010; Udeshi et al., 2013). Such proteomic approaches are particularly effective to globally analyse the ubiquitylome but are costly and burdensome to investigate the ubiquitylation of one or a selected set of proteins. Multiple assays have been devised to monitor and quantify ubiquitylation reactions performed *in vitro* using recombinant proteins or extracts (Berndsen and Wolberger, 2011; Boisclair et al., 2000; Chong et al., 2019; Gururaja et al., 2005; Mondal et al., 2016; Schneider et al., 2012), but they do not permit to investigate ubiquitylation events that take place in cells or tissues. This is generally achieved using conventional band shift assays and immunoblotting methods, often after an affinity purification of the ubiquitylated proteins (Hershko et al., 1982; Hjerpe et al., 2009; Hovsepian et al., 2016; Kaiser and Tagwerker, 2005). However, due to the low stoichiometry and the unstable nature of many ubiquitin conjugates, these methods are often not sufficiently sensitive to robustly assay the modification of endogenous proteins. Numerous studies thus rely upon overexpression of ubiquitin and/or substrate proteins, which may subvert endogenous ubiquitin conjugation pathways. There is therefore a need for alternative sensitive methods to probe the ubiquitylation of endogenously expressed proteins.

Protein-fragment complementation assays (PCAs) are a family of techniques devised to probe the proximity of proteins (Michnick et al., 2007). They rely on the use of complementary fragments of a reporter that are genetically fused to proteins of interest. These fragments can reconstitute the active reporter when brought into close proximity through the association of their fusion partners. The activity of the reporter is thus an indirect measure of the association of proteins fused to the reporter fragments. We reasoned that although generally used to probe non covalent protein interactions, PCAs could also be utilized to demonstrate the conjugation of ubiquitin to its substrate proteins. In this manuscript, we describe NUbiCA, a PCA based on the NanoLuc luciferase designed to probe the ubiquitylation of select proteins. Using budding yeast as a model system, we show that NUbiCA is a sensitive method that enables to examine mono- or poly-ubiquitin signals conjugated to proteins expressed at endogenous levels.

## RESULTS

### Design of a NanoLuc-based ubiquitin conjugation assay (NUbiCA)

NanoLuc is one of the smallest and brightest luciferase currently available (Hall et al., 2012). It has previously been further engineered to develop NanoBiT, a protein complementation reporter consisting of two asymmetricaly sized fragments (Dixon et al., 2016). The large fragment, termed LgBiT, retains an independent tertiary structure and has been optimized to have a high thermal stability and slow turnover. The small fragment, termed SmBiT, is a 11 amino acid peptide that has been selected for its low intrinsic affinity for LgBiT (their dissociation constant is ∼200 μM), while retaining the ability to reconstitute a bright luciferase. These properties make NanoBiT an excellent reporter to establish a ubiquitin conjugation assay, by fusing SmBiT to ubiquitin and LgBiT to ubiquitylation substrates. Ubiquitylation of the substrate places SmBiT in proximity to LgBiT, allowing the reassembly of an active luciferase (Figure 1a). SmBiT needs to be fused at the N-terminus of ubiquitin to preserve the C-terminal carboxyl group required for ubiquitin attachment to its substrates. In contrast, LgBiT can be positioned at either extremities of substrate proteins, or even in an internal loop (see Cse4 example hereafter). To ensure that the luminescence signal originates from the conjugation of SmBiT-ubiquitin to the LgBiT-tagged proteins and not to one of their interaction partners, we appended a polyhistidine tag to the LgBiT sequence. This enables to purify the ubiquitylation substrates under fully denaturing conditions before measuring the luminescence signal in the eluate (Figure 1b). Importantly, the total amount of purified LgBiT/His tagged proteins can readily be quantified using a high affinity SmBiT peptide variant termed HiBiT, which binds LgBiT with nanomolar affinity (Schwinn et al., 2018a). For control purposes, it is also needed to evaluate the expression level of SmBiT-ubiquitin. This can be done by measuring the luminescence of total protein extracts in the presence of recombinant LgBiT (Figure 1b). Altogether, the ubiquitylation level of LgBiT/His tagged proteins can be compared in different conditions using a normalized luminescence ratio (NLR, Figure 1c).

**Figure 1.**
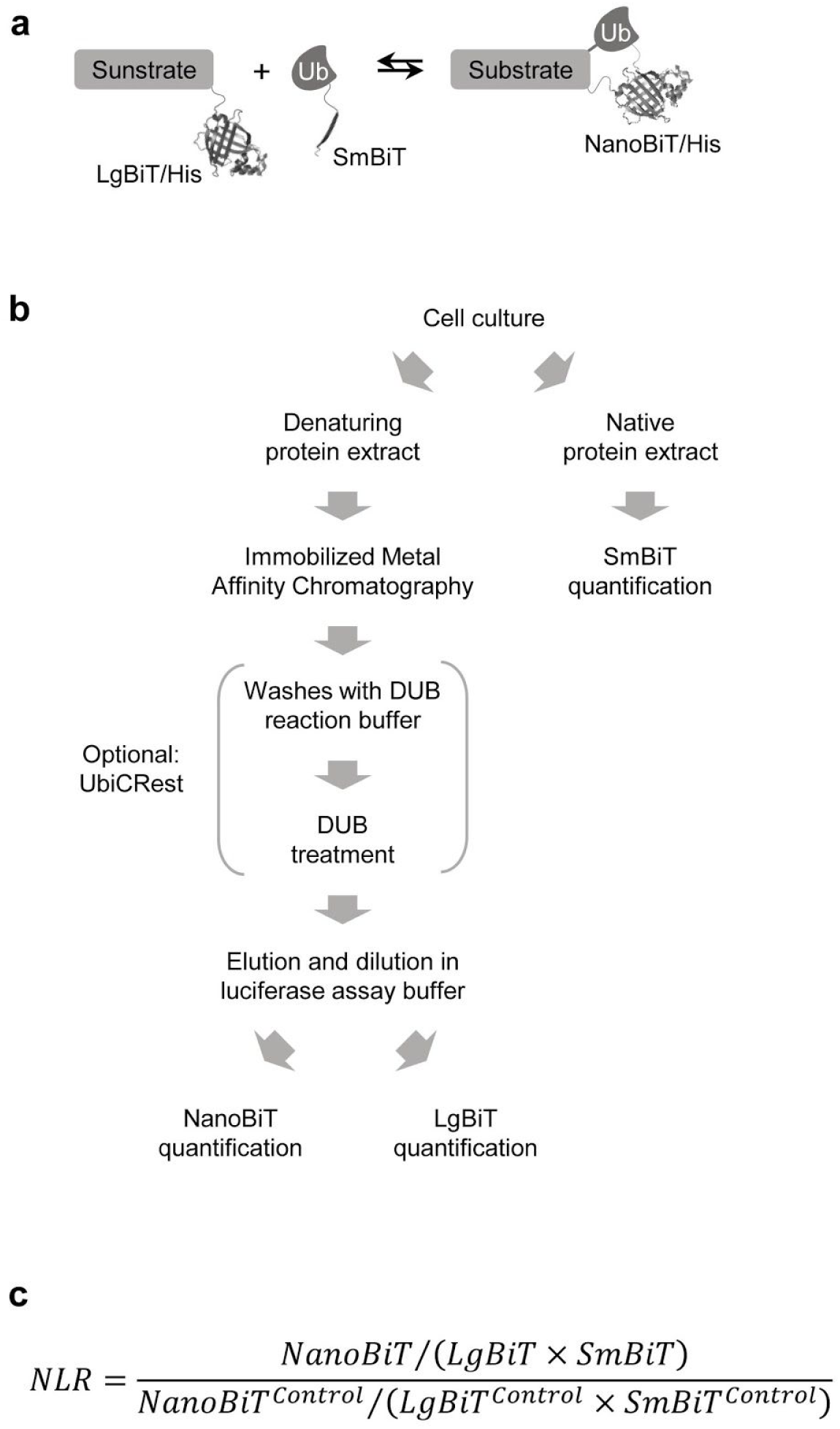
Description of the NanoLuc-based Ubiquitin Conjugation Assay (NUbiCA). (**a**) Ubiquitin (ub) and a substrate protein of interest are genetically fused to the LgBiT/His and SmBiT tags, respectively. Conjugation of ubiquitin to the protein of interest enables the reconstitution of an active luciferase. (**b**) Cells are lysed in denaturing and native conditions. The denaturing extract is used to pull down the LgBiT/His-tagged substrate with Immobilized Metal Affinity Chromatography. The resin is washed to remove the SmBiT-ubiquitin that is not conjugated to the substrate. For UbiCREst analysis, the resin is further washed with a DUB reaction buffer to ensure proper renaturation of ubiquitin conjugates before DUB digestion. The purified substrate is eluted and then diluted in a luciferase assay buffer. The luminescence of the eluate is measured either with or without the HiBiT peptide that tightly binds LgBiT and reconstitutes an active luciferase (Schwinn et al., 2018a). The luminescence measured without the HiBiT peptide enables to quantify the amount of ubiquitin-conjugated substrate present in the eluate (NanoBiT quantification). The luminescence measured with the HiBiT peptide enables to quantify the total amount of LgBiT/His-tagged substrate present in the eluate (LgBiT quantification). The native protein extract is supplemented with recombinant LgBiT to quantify the expression level of SmBiT-ubiquitin (SmBiT quantification). (**c**) The relative ubiquitylation level of the substrate is expressed as a normalized luminescence ratio (NLR). In each experiment, the NanoBiT, LgBiT and SmBiT signals are quantified in control and test conditions (e.g. wild type and mutant cells). LgBiT and SmBiT signals are used to correct NanoBiT signals for possible variations of substrate or SmBiT-ubiquitin levels (see also Supplementary Note 1). The NLR corresponds to the corrected NanoBiT signal of a test condition normalized by the corrected NanoBiT signal of the control condition.

### Validation of NUbiCA to probe proteolytic and non-proteolytic ubiquitylation events

We used budding yeast as a model organism to evaluate NUbiCA. We first probed the ubiquitylation of H2B, one of the best characterized ubiquitin conjugate in cells. In yeast, H2B is mono-ubiquitylated on its lysine K124 by the Bre1 ubiquitin ligase and the Rad6 conjugating enzyme Rad6 (Hwang et al., 2003; Robzyk et al., 2000; Wood et al., 2003) (Figure 2a). We endogenously fused *HTB2*, one of the two H2B producing genes in yeast, with a C-terminal LgBiT/His tag and constructed wild type and mutant strains expressing both SmBiT-ubiquitin and Htb2-LgBiT/His. Using a classical band shift assay, we verified that SmBiT-ubiquitin can be efficiently conjugated to Htb2 in wild type cells, but not in *bre1*Δ, *rad6*Δ or *htb2(K124R)* mutants (Figure 2b). This shows that the SmBiT and the LgBiT/His tags do not significantly perturb the ubiquitylation of Htb2. We then assessed Htb2 ubiquitylation using NUbiCA. We observed that Htb2-LgBiT/His purified from wild type cells produced a robust luminescent signal, which was largely reduced in *bre1*Δ, *rad6*Δ and *htb2(K124R)* mutants (Figure 2c). Thus, the results obtained with NUbiCA correlate with those obtained with a band shift assay, indicating that NUbiCA accurately reports Htb2 mono-ubiquitylation.

**Figure 2.**
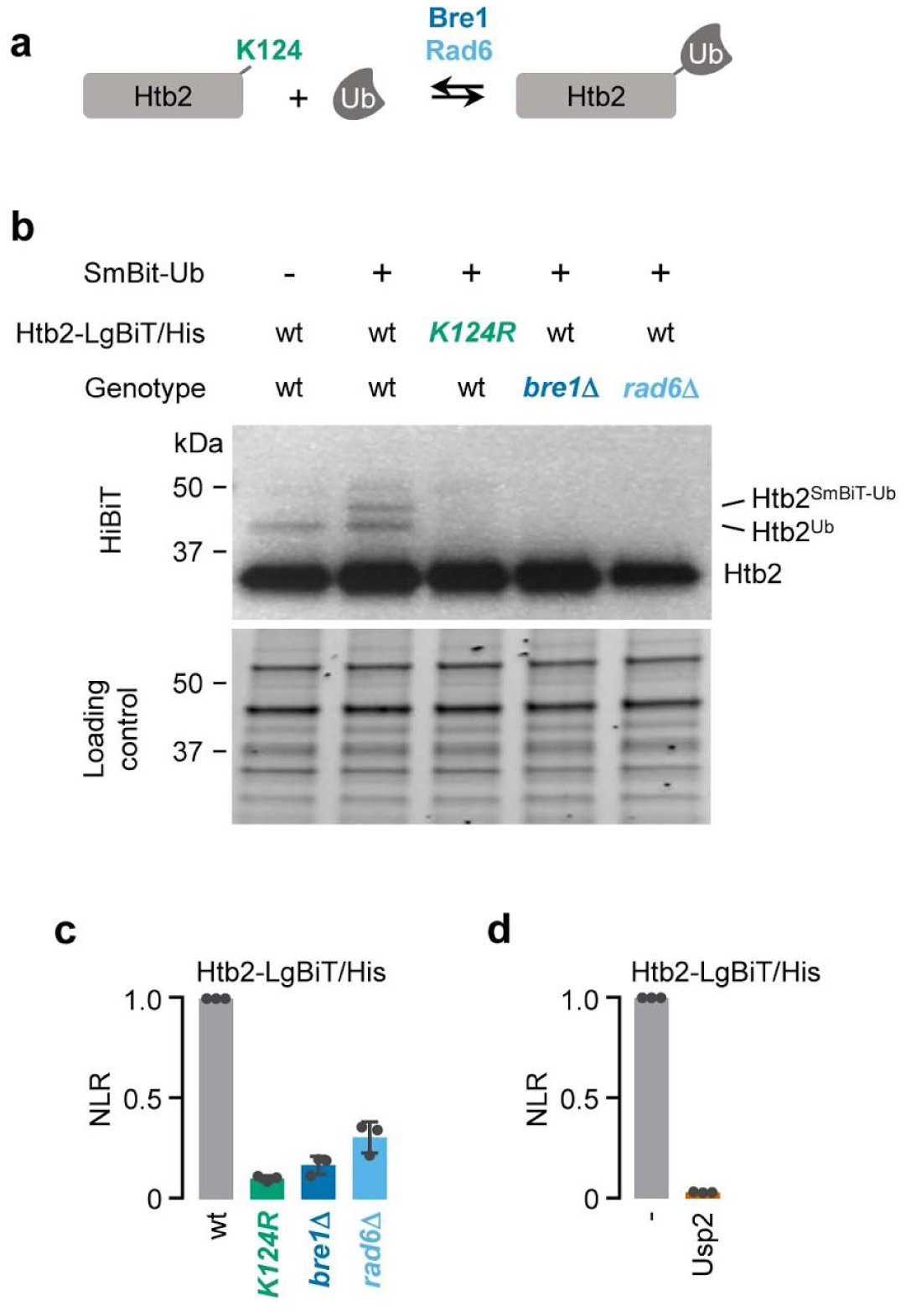
Analysis of Htb2 mono-ubiquitylation. (**a**) Htb2 is ubiquitylated on its lysine K124 by the Bre1 ubiquitin ligase and the Rad6 ubiquitin conjugating enzyme. (**b**) SmBiT-Ubiquitin is conjugated to Htb2-LgBiT/His in wild type cells, but not in *htb2(K124R), bre1*Δ or *rad6*Δ, or mutants. Total protein extracts (prepared from the indicated yeast strains expressing LgBiT/His-tagged Htb2) were separated by SDS-PAGE. Proteins loaded on the gel were revealed using Stain-Free™ imaging (Loading control) and transferred onto a nitrocellulose membrane. Htb2-LgBiT/His was visualised with a gel imager after incubating the membrane with the HiBiT peptide, which tightly interacts with LgBiT to reconstitute a functional luciferase (HiBiT). (**c**) Relative ubiquitylation levels of Htb2-LgBiT/His purified from the indicated yeast strains. The graph displays normalized luminescence ratios (NLR) (mean ± s.d., n=3). (**d**) Ubiquitin Chain Restriction analysis of Htb2-LgBiT/His. Htb2-LgBiT/His was purified from wild type cells and treated with Usp2. The graph displays normalized luminescence ratios (NLR) (mean ± s.d., n=3).

Histone mono-ubiquitylation acts non-proteolytically to control gene activity (Weake and Workman, 2008). However, proteins modified by poly-ubiquitin chains or multiple mono-ubiquitin are often rapidly degraded by the proteasome. Such proteolytic ubiquitylation events can be difficult to assay without using artificial conditions such as overexpression or proteasome inhibition. We therefore wished to test whether NUbiCA enables to detect the ubiquitylation of unstable proteins in endogenous conditions. We chose to probe the ubiquitylation of the N-terminal region of the transcription factor Stp2 (Stp2^N^), which comprises a strong degradation signal (Omnus and Ljungdahl, 2014). Using proteasome inhibition and classical immunoassays, we previously demonstrated that Stp2^N^ is efficiently ubiquitylated, primarily by the Asi1/3 ubiquitin ligase complex and the Ubc7 conjugating enzyme (Khmelinskii et al., 2014) (Figure 3a). Using NUbiCA, we observed that endogenously expressed Stp2^N^-LgBiT/His purified from wild type cells expressing SmBiT-ubiquitin produced a clear luminescence signal, which was decreased in *asi3*Δ and *ubc7*Δ mutants (Figure 3b). This indicates that NUbiCA is able to reveal the modification of a short-lived ubiquitylated protein expressed at endogenous levels without proteasome inhibition.

**Figure 3.**
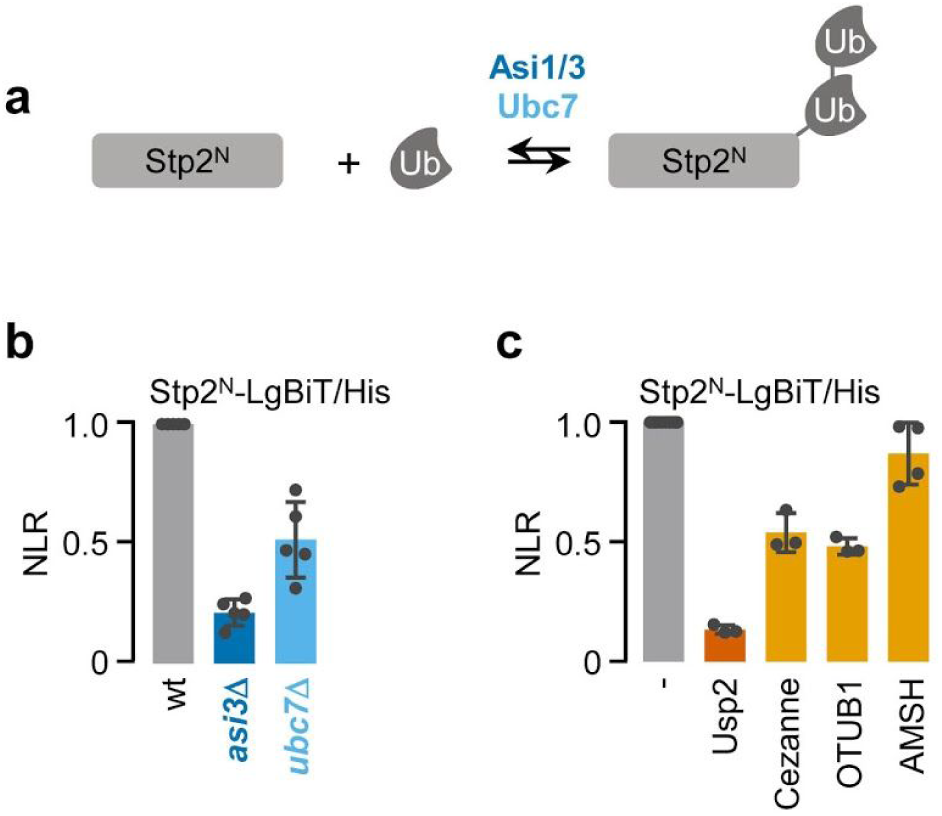
Analysis of Stp2 poly-ubiquitylation. (**a**) Stp2 is primarily ubiquitylated by the Asi1/3 ubiquitin ligase complex and the Ubc7 ubiquitin conjugating enzyme. (**b**) Relative ubiquitylation levels of Stp2^N^-LgBiT/His purified from the indicated yeast strains. The graph displays normalized luminescence ratios (NLR) (mean ± s.d., n=5). (**c**) Ubiquitin Chain Restriction analysis of Stp2^N^-LgBiT/His. Stp2^N^-LgBiT/His was purified from wild type cells and treated with the indicated DUBs. The graph displays normalized luminescence ratios (NLR) (mean ± s.d., n≥3).

### NUbiCA can be combined with UbiCRest to dissect ubiquitin chain linkages

One of the most important challenges in the field of protein ubiquitylation is to decipher how ubiquitin signals regulate the fate and activity of cellular proteins. Multiple tools and methods have been devised to investigate ubiquitin chain topologies, including ubiquitin mutants, linkage-specific reagents and mass spectrometry (Hospenthal et al., 2015; Mattern et al., 2019; Meyer and Rape, 2014; Newton et al., 2008; Ordureau et al., 2015; Spence et al., 1995). We thought to combine NUbiCA with UbiCRest, a method that uses linkage specific deubiquitylation enzymes (DUBs) to dissect ubiquitin chains (Hospenthal et al., 2015). Since DUBs recognize the native ubiquitin fold, purified LgBiT/His proteins need to be refolded before incubation with DUBs (Figure 1b). To evaluate the refolding efficiency, we incubated purified Htb2-LgBiT/His and Stp2^N^-LgBiT/His with Usp2, a non-specific DUB that hydrolyses all types of ubiquitin linkages (Hospenthal et al., 2015). This resulted in a large reduction of the luminescence signals produced by both proteins (Figures 2d and 3c), showing that ubiquitin is indeed efficiently refolded in our experimental conditions. Next, we incubated purified Stp2^N^-LgBiT/His with Cezanne, OTUB1 and AMSH, 3 DUBs that preferentially hydrolyse K11-, K48- and K63-linked ubiquitin chains, respectively (Hospenthal et al., 2015). The DUBs concentrations and incubation time were adjusted to ensure their specificity (Supplementary Figure 1). Incubation of Stp2^N^-LgBiT/His with Cezanne and OTUB1 reduced the signal approximately 2 fold, while AMSH had only a marginal effect (Figure 3c). This indicates that Stp2^N^-LgBiT/His is modified with poly-ubiquitin chains comprising proteolytic K11 and K48 linkages. This result is consistent with the role of the Asi1/3 complex in the degradation and negative regulation of Stp2 activity (Boban et al., 2006; Khmelinskii et al., 2014).

### Ubiquitylation of the centromeric histone Cse4

In order to further test the ability of NUbiCA to dissect ubiquitin signalling, we chose to investigate the ubiquitylation of the yeast histone variant Cse4. Cse4 is an essential protein that substitutes for histone H3 in centromeric nucleosomes (Meluh et al., 1998). With ∼100 copies per cell, Cse4 is among the 20% least expressed proteins in yeast (Kulak et al., 2014). When overexpressed, Cse4 ectopically localizes to euchromatin, which perturbs transcription and causes chromosome instability (Au et al., 2013; Hildebrand and Biggins, 2016). The ubiquitin ligase Psh1 is described to control Cse4 levels and to prevent the mislocalization of overexpressed Cse4 (Hewawasam et al., 2010; Ranjitkar et al., 2010) (Figure 4a), but whether and when it ubiquitylates endogenous Cse4 has not been directly demonstrated.

**Figure 4.**
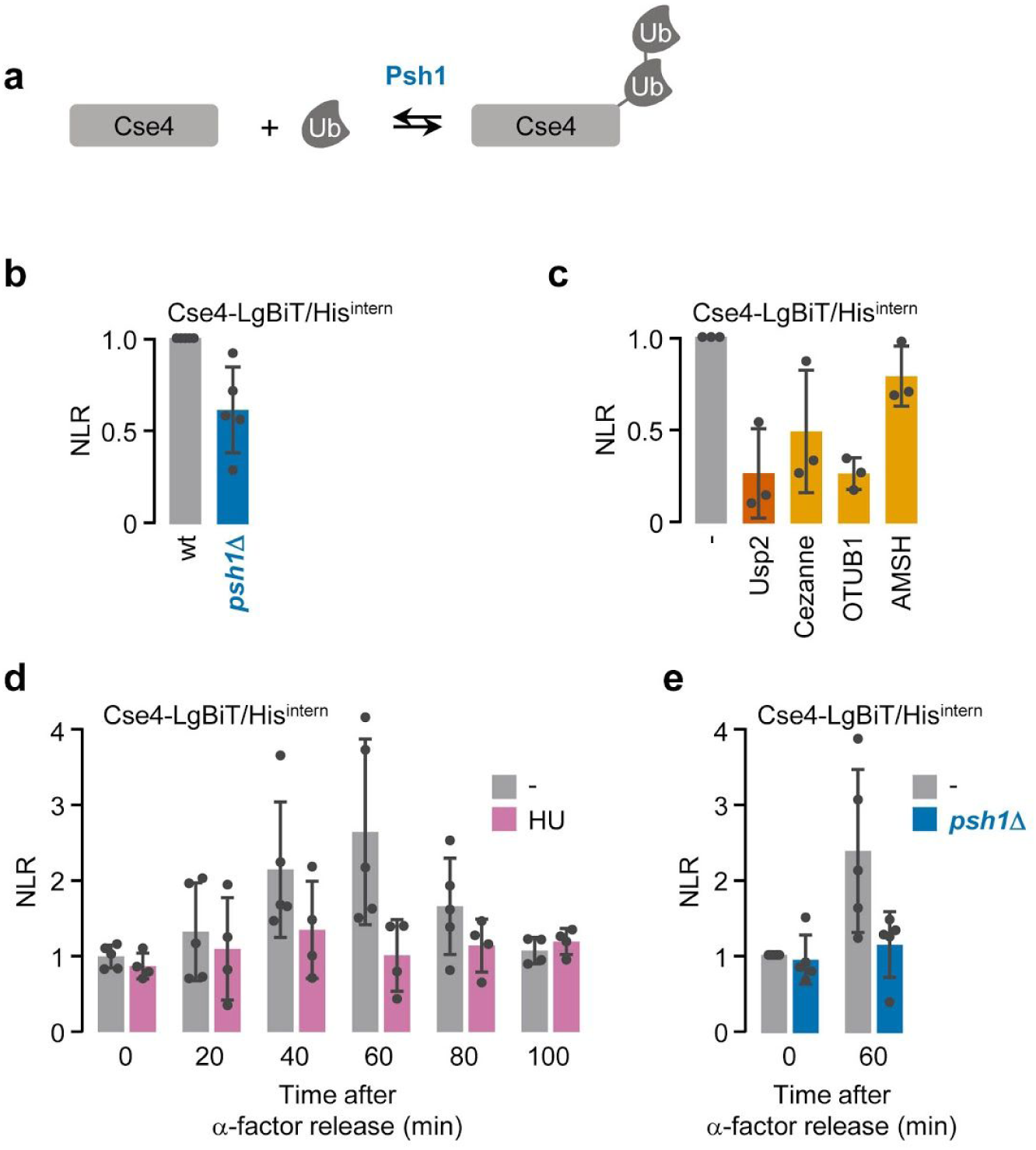
Analysis of Cse4 ubiquitylation. (**a**) Cse4 is ubiquitylated by the Psh1 ubiquitin ligase. (**b**) Relative ubiquitylation levels of Cse4-(LgBiT/His)^intern^ purified from wild type (wt) and *psh1*Δ cells. The graph displays normalized luminescence ratios (NLR) (mean ± s.d., n=5). (**c**) Ubiquitin Chain Restriction analysis of Cse4-(LgBiT/His)^intern^. Cse4-(LgBiT/His)^intern^ was purified from wild type cells and treated with the indicated DUBs. The graph displays normalized luminescence ratios (NLR) (mean ± s.d., n=3). (**d**) Relative ubiquitylation levels of Cse4-(LgBiT/His)^intern^ after α-factor release. Wild type cells expressing Cse4-(LgBiT/His)^intern^ were synchronized in G1 with α-factor, released in the absence (grey) or presence of hydroxyurea (HU, pink) and sampled at the indicated timepoints. The graph displays normalized luminescence ratios (NLR) (mean ± s.d., n≥4). (**e**) Relative ubiquitylation levels of Cse4-(LgBiT/His)^intern^ in wild type and *psh1*Δ cells during S-phase. Cse4-(LgBiT/His)^intern^ was purified from wild type (grey) and *psh1*Δ (blue) cells synchronized in G1 with α-factor and sampled at 60 minutes after release. The graph displays normalized luminescence ratios (NLR) (mean ± s.d., n=5).

To address this question, we tagged endogenous Cse4 with LgBiT/His in cells expressing SmBiT-ubiquitin. The LgBiT/His tag was inserted at an internal position (between asparagine 80 and leucine 81), as this was shown to minimally perturb Cse4 function (Wisniewski et al., 2014). When we purified Cse4-(LgBiT/His)^intern^ from wild type cells, we could detect a weak luminescence signal, which decreased in a *psh1*Δ mutant (Figure 4b). This suggests that Psh1 can ubiquitylate Cse4 expressed at endogenous levels. UbiCRest analysis showed reduced luminescence after incubation with Usp2, Cezanne and OTUB1 (Figure 4c), indicating that Cse4 is modified by ubiquitin chains containing K11 and K48 linkages.

During S-phase, the pre-existing pool of Cse4 is replaced by newly synthetized Cse4 (Wisniewski et al., 2014). To determine whether this is accompanied by ubiquitin dependent regulation, we synchronized cells in G1 with α-factor and used NUbiCA to monitor Cse4 ubiquitylation at various time points after release. We observed a peak of ubiquitylation 60 minutes after release (Figure 4d), which corresponds to the time at which centromeric Cse4 is exchanged (Wisniewski et al., 2014). This peak was not detected when the cells were treated with hydroxyurea to block DNA replication (Figure 4d) or in *psh1*Δ cells (Figure 4e), although these cells progressed normally through the cell cycle (Supplementary Figure 2). Altogether, these results suggest that Psh1-dependent ubiquitylation regulates the degradation of endogenous Cse4 during DNA replication. We propose that this ubiquitylation could serve to promote the replacement of centromeric Cse4 and/or to correct erroneous incorporation of newly synthetized Cse4 in euchromatin. Further investigation will be required to address this question as well as the possible role of other ubiquitin ligases that have been described to regulate overexpressed Cse4 (Cheng et al., 2016; Cheng et al., 2017; Ohkuni et al., 2016).

### A genome wide collection of yeast strains ready for NUbiCA

To facilitate the investigation of protein ubiquitylation using NUbiCA, we thought to construct a genome wide collection of yeast strains expressing SmBiT-ubiquitin and proteins C-terminally tagged with LgBiT/His. We used the recently established C-SWAT yeast library (Meurer et al., 2018) to systematically fuse yeast open reading frames to the DNA sequence of the LgBIT/His tag. The resulting strains were then crossed with a strain containing a chromosomally integrated SmBiT-ubiquitin expression cassette. The entire procedure was successful for more than 98% of the colonies from the original C-SWAT collection, yielding a collection of 5,580 ready-to-use NUbiCA strains (Supplementary Table 1).

## DISCUSSION

The methods most commonly used to demonstrate the ubiquitylation of a protein of interest rely on the separation of the ubiquitylated and unmodified protein forms by gel electrophoresis. However, these methods are not always sufficiently sensitive to detect the ubiquitylation of proteins expressed at endogenous levels. They are also difficult to quantify and not well amenable to large scale studies. Recently, alternative assays based on ELISA and FRET approaches been established to quantify ubiquitylated proteins in cell or tissue extracts (Foote et al., 2018; Guven et al., 2019). Because these assays have only been recently reported, it is difficult to evaluate their sensitivity and specificity for the detection of low abundance ubiquitylated proteins.

In the present study, we demonstrate that the conjugation of ubiquitin to proteins can also be monitored using a protein complementation assay. One of the limitations of this strategy is that the proteins of interest and ubiquitin have to be tagged with fragments of a complementation reporter. Hence this method is only applicable to study ubiquitylation in tissue culture systems or genetically amenable model organisms. We chose NanoBiT as a complementation reporter for two main reasons. First, the NanoBiT fragments, LgBiT and SmBiT, are small, stable and have very low intrinsic affinity (Dixon et al., 2016). They are thus less likely to perturb the function and the ubiquitylation of the proteins they are fused to. Indeed, it is known that ubiquitin N-terminally fused to small tags can functionally replace endogenous ubiquitin (Ling et al., 2000). Second, NanoLuc is one of the brightest luciferase, with a detection limit of less than 1 amol (Hall et al., 2012; Schwinn et al., 2018a). We reasoned that this exquisite sensitivity should enable the detection of scarce ubiquitylated proteins. Indeed, we could successfully detect proteolytic ubiquitylation of Stp2^N^ and of the low abundance histone Cse4 expressed from their endogenous chromosomal locus, suggesting that NUbiCA will be applicable to investigate the ubiquitylation of most cellular proteins without overexpression. This is in our view of utmost important since overexpression can trigger protein quality control and ubiquitylation pathways that are not necessarily the ones one wants to investigate. For instance, using endogenously expressed and internally tagged Cse4, we could show that Cse4 is specifically ubiquitylated during S-phase (Figure 4d). This finding could not have been made using overexpressed Cse4, which is massively ubiquitylated to prevent its accumulation in euchromatin (Ranjitkar et al., 2010).

An important aspect of the NUbiCA protocol described here is that the lysate preparation and substrate purification are done under highly denaturing conditions (Figure 1b). This has the advantage to suppress DUB activity and, therefore, to preserve ubiquitin conjugates. It also ensures that the luminescence signal originates from ubiquitin conjugated to the LgiT/His-tagged protein and not to one of its interaction partners. Thus, the denaturing purification is important to achieve a high sensitivity and specificity. Yet, it is possible that this step could be omitted for certain proteins. The ubiquitylation of these proteins could then be monitored directly in extracts or even in intact cells. Indeed, NanoBiT has recently been used to monitor the modification of Cullin1 by the ubiquitin like protein Nedd8 in mammalian cells (Schwinn et al., 2018b). It will be important to determine to what extent this approach can be applied to other proteins since it would open the possibility to monitor ubiquitylation and deubiquitylation reactions in real time in live cells.

Other important features of the NUbiCA protocol is that it is generic and does not involve gel electrophoresis. It will thus be easier to perform larger scale studies than with the currently available assays, for instance to compare the ubiquitylation of a protein of interest in multiple conditions, or to investigate the ubiquitylation of multiple proteins in parallel. Moreover, we demonstrate that NUbiCA is compatible with UbiCRest, a method previously established to distinguish ubiquitin linkages (Hospenthal et al., 2015). The genome wide collection of NUbiCA yeast strains that we constructed will thus be a powerful resource that will facilitate the investigation of the ubiquitin code. Finally, although we limited our study to ubiquitylation in yeast, the principles of NUbiCA can be generalized to monitor other posttranslational protein modifiers, such as SUMO or Nedd8, in any tissue culture system or genetically amenable model organism.

## MATERIALS AND METHODS

### Yeast methods and plasmids

Yeast genome manipulations (chromosomal gene tagging and gene deletion) were done using conventional procedures based on PCR targeting and plasmid integration. Cassettes for PCR targeting were amplified with the Phusion DNA polymerase (New England Biolabs). Gene deletions and tagging were validated by PCR. Yeast strains and plasmids used in this study are listed in Supplementary Tables 2 and 3, respectively.

### Purification of LgBiT/His-tagged proteins and UbiCRest

LgBiT/His-tagged proteins were purified from 10^8^ to 10^9^ exponentially growing yeast cells. Cell pellets were resuspended in 20% trichloroacetic acid and lysed with glass beads in a Disrupter Genie homogenizer (Scientific Industries). After precipitation, proteins were resuspended in a denaturing extraction buffer (6 M guanidinium chloride, 100 mM Tris-HCl, pH 9, 300 mM NaCl, 0.2% Triton X-100 and 5 mM chloroacetamide), clarified at 30,000 g and incubated for 90 min at room temperature with TALON® Metal Affinity Resin (Clontech). The beads were then washed twice with the extraction buffer and twice with a wash buffer (2 M urea, 100 mM sodium phosphate pH 7.0, 300 mM NaCl). LgBiT/His-tagged proteins were finally eluted with an elution buffer (2 M urea, 100 mM sodium phosphate pH 7.0, 300 mM NaCl, 250 mM imidazole).

In case of UbiCRest, the beads washed with the extraction buffer were further washed with a DUB reaction buffer (50 mM Tris pH 7.5, 100mM NaCl, 5 mM 2-mercaptoethanol). They were then incubated with recombinant DUBs at 37°C for 1 h (Usp2, Enzo Life Sciences) or overnight (GST-AMSH, GST-Cezanne or GST-OTUB1). After incubation, the beads were washed twice with the wash buffer and eluted with the elution buffer.

### Luminescence measurements

To measure the NanoBiT signal and quantify the amount of purified LgBiT/His-tagged substrates, the eluates were diluted in a luciferase assay buffer (PBS, 0.1% BSA) with or without the HiBiT peptide (0.5 µM, VSGWRLFKKIS, synthetized by ProteoGenix) and mixed with furimazine (0.25% Nano-Glo® Substrate, Promega). The HiBiT peptide binds tightly to LgBiT (K_D_ = 0.7 nM) and reconstitutes an active luciferase (Schwinn et al., 2018a). The samples were distributed in duplicates in 96 half-well white polystyrene microtiter plates and spaced with blank wells to avoid signal crosstalk from the HiBiT containing samples. Luminescence was measured with either a Xenius XL luminometer (SAFAS) or a FLUOstar Omega plate reader (BMG labtech).

To measure the expression level of SmBiT-ubiquitin, cell pellets were resuspended in PBS and lysed with glass beads in a Disrupter Genie homogenizer. After clarification, the extracts were mixed with recombinant GST-LgBiT (0.05 µM) and furimazine (0.25% Nano-Glo® Substrate, Promega) and distributed in microtiter plates for luminescence measurements.

### Recombinant protein expression and purification

*E. coli* BL21(DE3)RIL cells were transformed with plasmids encoding GST-AMSH (pGEX4T2_GST-AMSH), GST-OTUB1 (pOPINK_OTUB1), GST-Cezanne (pOPINK_Cezanne) or GST-LgBiT (pGR0890) and were cultivated in LB medium. Protein expression was induced by addition of 1 mM IPTG during 4 h at 23.5°C. Cells were pelleted, resuspended in a lysis buffer (PBS, 0.05% lysozyme, 1 mM DTT), and lysed by sonication. Lysates were rotated with glutathione beads (GE Healthcare) for 45 min at 4°C. Beads were washed, first with PBS, 1 mM DTT, then with 50 mM Tris pH 7.8, 200 mM NaCl, 5 mM DTT. Purified proteins were eluted in 50 mM Tris pH 7.8, 200 mM NaCl, 5 mM DTT, 20mM reduced L-Glutathione and dialyzed against PBS, 10% glycerol, 1 mM DTT. Protein purity was tested using Stain-Free™ imaging (Bio-Rad). Protein concentration was estimated by absorbance at 280 nm.

### DUB specificity assays

K11-, K48- and K63-linked tetra-ubiquitin chains (Boston Biochem) were diluted to a concentration of 3 µM in 50 mM HEPES pH 7.5, 50 mM NaCl and incubated overnight at 37°C with recombinant DUBs. The digested ubiquitin chains were then denatured in Laemmli sample buffer, separated by SDS-PAGE and detected with silver staining.

### Western blots

Total protein extracts were prepared from 2.10^8^ exponentially growing yeast cells expressing Htb2-LgBiT/His. Cell pellets were resuspended in 20% trichloroacetic acid and lysed with glass beads in a Disrupter Genie homogenizer (Scientific Industries). After precipitation, proteins were solubilized in TCA sample buffer (450 mM Tris pH 8.8, 1% SDS, 2 mM EDTA, 15% glycerol, 100 mM DTT, Bromophenol blue). Protein samples were clarified, denatured and analyzed by SDS–PAGE using 4-20% Mini-PROTEAN® TGX Stain-Free™ precast gels (Bio-Rad). Total protein levels were assessed Stain-Free™ imaging (Bio-Rad). Proteins were then transferred on a nitrocellulose membrane with a Trans-Blot® Turbo™ semi-dry transfer apparatus (Bio-Rad). To reveal LgBiT/His-tagged Htb2, the membrane was blocked with TBS-T (50 mM Tris-Cl pH 7.5, 150 mM NaCl, 0.05% TWEEN® 20) and incubated for 5 minutes with TBS-T supplemented with 1 µM HiBiT peptide (VSGWRLFKKIS, synthetized by ProteoGenix) and 0.4% Nano-Glo® Substrate (Promega). Luminescence was detected using an Amersham Imager 680.

### Cell cycle synchronization and analysis

Overnight yeast cultures were diluted to an OD_600_ of 0.2 and grown for 3 h in YPD medium. Cells were then treated with α-factor (5 μg.mL^-1^, GenScript) for 90 min, collected by filtration, washed with PBS and released into YPD medium with or without hydroxyurea (15 mg.mL^-1^, Euromedex).

For cell cycle analysis, culture samples were fixed with 75% ethanol and treated for 1 h at 37°C with RNase (1 mg.mL^-1^ in 10 mM Tris-HCl, 15 mM NaCl) and then for 1 h at 50°C with Proteinase K (1 mg.mL^-1^ in PBS). After sonication, cells were stained with propidium iodide (0.6 μg.mL^-1^). DNA content was analysed with a FACSAria™ II flow cytometer (BD Biosciences) using a 488 nm excitation laser and a 616/23 bandpass filter. 10,000 events were collected for each condition.

### Construction of a collection of NUbiCA yeast strains

The SWAP-Tag approach was used to assemble a genome wide collection of yeast strains coexpressing SmBiT-ubiquitin and proteins C-terminally tagged with LgBiT/His. Meurer et al. previously constructed a library of 5,661 yeast strains where an acceptor module has been integrated before the stop codon of individual ORFs (Meurer et al., 2018). This C-SWAT acceptor module can be efficiently exchanged with a donor module provided by a plasmid. The plasmid pAB0010 (which provides a donor module containing a LgBiT/His tag followed by a heterologous terminator and a truncated Hygromycine B resistance marker) was transformed into the yMAM1205 strain. The transformed strain was crossed with the full collection of C-SWAT strains arrayed in a 384-colony format using a ROTOR HDA pinning robot (Singer Instruments). The colonies were sequentially pinned on appropriate media to select diploids, sporulate, select haploids, induce the exchange of the acceptor and donor modules and select the recombinant strains using the procedure described by Meurer et al. (Meurer et al., 2018). This produced a collection a MAT-α haploid strains expressing proteins C-terminally tagged with LgBiT/His. This collection was then crossed with the scGLD0122 strain, which contains an SmBiT-ubiquitin expression cassette inserted at the *MET17* locus flanked with a Nourseothricin resistance marker. Again, the colonies were pinned on appropriate selection media to finally obtain MAT-α haploid strains coexpressing SmBiT-ubiquitin and proteins C-terminally tagged with LgBiT/His. The entire procedure was successful for more than 98% of the colonies, yielding a collection of 5,580 NUbiCA strains.

## ACKNOWLEDGEMENTS

We thank Matthias Meurer and Michael Knop for discussions, reagent exchanges, the C-SWAT library and their help with the construction of the NUbiCA yeast strain collection. We also thank David Komander and Sylvie Urbe for DUB expression plasmids, Rémy Le Guevel for the ImPACcell platform and the IGDR and Biosit for support and infrastructure.

## AUTHOR CONTRIBUTIONS

G.R. coordinated the project and designed the experiments. M.L.B. and A.B performed the NanoBiT experiments. G.L.D and A.B. constructed the NUbiCA yeast strain collection. G.R. and M.L.B. prepared the figures and wrote the manuscript.

